# Unmet Needs for Analyzing Biological Big Data: A Survey of 704 NSF Principal Investigators

**DOI:** 10.1101/108555

**Authors:** Lindsay Barone, Jason Williams, David Micklos

## Abstract

In a 2016 survey of 704 National Science Foundation (NSF) Biological Sciences Directorate principal investigators (BIO PIs), nearly 90% indicated they are currently or will soon be analyzing large data sets. BIO PIs considered a range of computational needs important to their work—including high performance computing (HPC), bioinformatics support, multi-step workflows, updated analysis software, and the ability to store, share, and publish data. Previous studies in the United States and Canada emphasized infrastructure needs. However, BIO PIs said the most pressing unmet needs are training in data integration, data management, and scaling analyses for HPC—acknowledging that data science skills will be required to build a deeper understanding of life. This portends a growing data knowledge gap in biology and challenges institutions and funding agencies to redouble their support for computational training in biology.

## Background

Genotypic data based on DNA and RNA sequence have been the major driver of biology’s evolution into a data science. The current Illumina HiSeq X sequencing platform can generate 900 billion nucleotides of raw DNA sequence in under three days—four times the number of annotated nucleotides currently stored in GenBank, the U.S. “reference library” of DNA sequences (*1, 2*). In the last decade, a 50,000-fold reduction in the cost of DNA sequencing (*3*) has led to an accumulation of 9.3 quadrillion (million billion) nucleotides of raw sequence data in the National Center for Biotechnology Information (NCBI) Sequence Read Archive (SRA). The amount of sequence in the SRA doubled on average every 6–8 months from 2007–16 (4). It is estimated that by 2025, the storage of human genomes alone will require 2–40 exabytes (*5*), where an exabyte of storage would hold 100,000 times the printed materials of the U.S. Library of Congress (*6*). Beyond genotypic data, Big Data are flooding biology from all quarters—phenotypic data from agricultural field trials, patient medical records, and clinical trials; image data from microscopy, medical scanning, and museum specimens; interaction data from biochemical, cellular, physiological, and ecological systems—as well as an influx of data from translational fields such as bioengineering, materials science, and biogeography.

A 2003 report of an NSF blue-ribbon panel, headed by Daniel Atkins, popularized the term *cyberinfrastructure* to describe systems of data storage, software, HPC, and people that can solve scientific problems of the size and scope presented by big data (*7*). The Atkins Report was the impetus for several cyberinfrastructure projects in the biological sciences—including the NSF’s CyVerse, the Department of Energy’s KBase, and the European Union’s ELIXIR. The report described cyberinfrastructure as the means to harness the data revolution and to develop a “knowledge economy.” Although people were acknowledged as active elements of cyberinfrastructure, few published studies have assessed how well their computational and cyberinfrastructure needs are being met.

In 2006, EDUCAUSE surveyed 328 information technology (IT) professionals, primarily chief information officers, at institutions in the U.S. and Canada (*8*). When asked about preferences for funding allocation, respondents rated training and consulting (20%) a distant second to infrastructure and storage (46%). This suggested that “training and consulting get short shrift when bumped against the realities of running an IT operation.” Infrastructure and training emerged as similarly important in a study done as part of the 2015 University of Illinois’ “Year of Cyberinfrastructure” (*9*). Faculty and graduate students responding to a survey (n = 327) said they needed better access to data storage (36%), data visualization (29%), and HPC (19%). Although it was not directly addressed in the survey, training—on a range of technologies and across skills levels—emerged as a major need in focus groups (n = 200).

Over the last four years, CyVerse has taken the computational pulse of the biological sciences by surveying attendees at major professional meetings. Consistently, and across different conference audiences, 94% of students, faculty, and researchers said that they currently use large data sets in their research or think they will in the near future (n = 1,097). Even so, 47% rated their bioinformatics skill level as “beginner,” 35% rated themselves “intermediate,” and 6% said they have never used bioinformatics tools; only 12% rate themselves “advanced” (n = 608). Fifty-eight percent felt their institutions do not provide all the computational resources needed for their research (n = 1,024). These studies suggest a scenario of big data inundating unprepared biologists.

## Methods

In summer 2016 we expanded upon our previous studies with a purposive needs assessment of principal investigators (PIs) with grants from the National Science Foundation Directorate of Biological Sciences (BIO). Working from a list of 5,197 active grant awards, we removed duplicate PIs and those without email addresses to produce a final list of 3,987 subjects. The survey was administered in Survey Monkey using established methods (*10*). An initial email invitation with a link to the survey was sent to each subject in June 2016, with three follow-up emails sent at two-week intervals. Surveys were completed by 704 PIs, a response rate of 17.7% that provided a +/-3.35% margin of error at the 95% confidence level.

The respondents were asked to consider 13 computational elements of research—including data storage, discovery, analysis, and sharing. For each need, PIs were asked to reflect on their current use, their anticipated future requirements, and the institutional resources available to meet the need. Data were analyzed in IBM SPSS Statistics version 23. “I don’t know” responses were eliminated from the analysis of computational needs questions. Frequencies were calculated for each of the affirmative and negative responses in the computational needs matrix. Chi-square tests for independence were used to determine if there were significant differences in computational needs across three dimensions: 1) NSF BIO division, 2) research area (bioinformatics/computational biology *versus* all others), and 3) research group size (groups of less than five *versus* groups with more than five).

## Results

The respondents were relatively evenly dispersed among four major BIO divisions: Division of Biological Infrastructure (DBI), Division of Environmental Biology (DEB), Division of Integrative Organismal Systems (IOS), and Division of Molecular and Cellular Biosciences (MCB). These BIO PIs worked with a variety of data—with sequence, image, phenotype, and ecological data predominating (Figure 1). The vast majority (87%) said they are currently using big datasets in their research or will within the next three years. This is slightly lower than in our previous studies of meeting attendees, a large proportion of whom had a genomics focus or were students or early career researchers.

**Figure 1.**
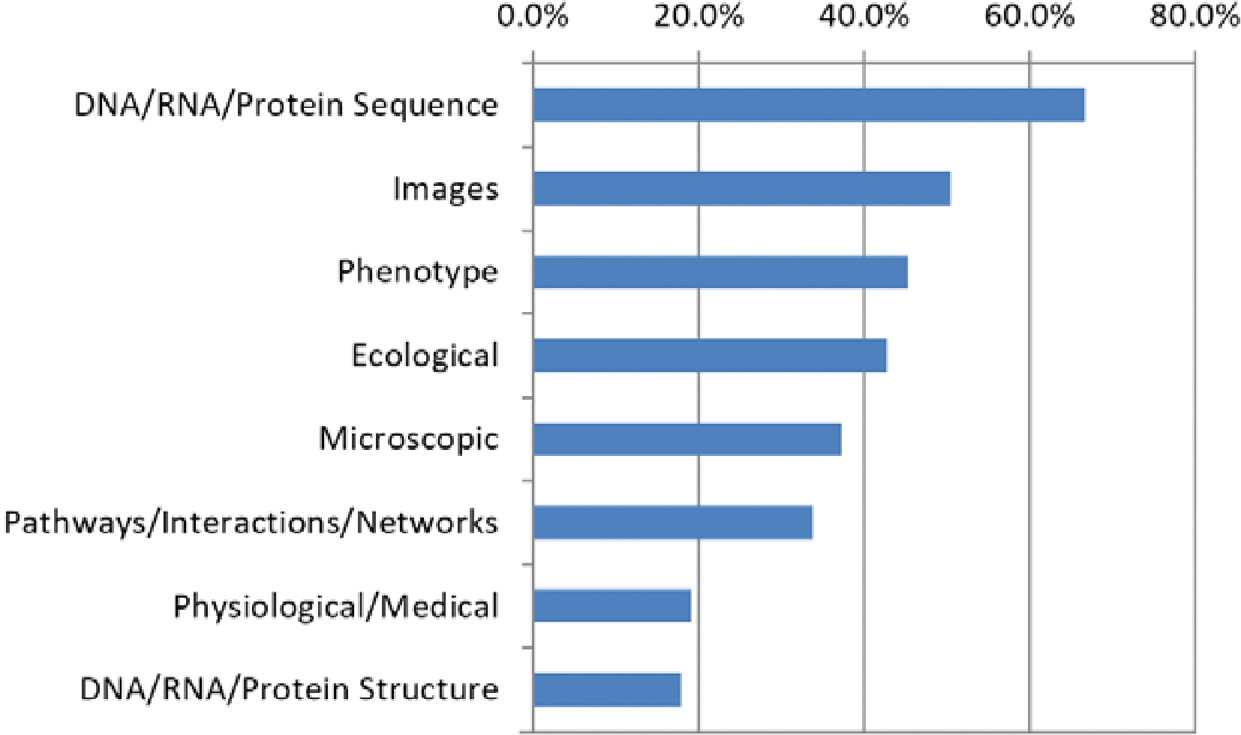
Major data types used by BIO PIs.

More than half of the PIs said that 11 of the 13 computational needs are currently important to their research. The proportions increased *across all needs*—to 82–97%—when PIs considered what would be important three years in the future (Figure 2). Significantly more PIs who identified themselves as bioinformaticians said nine of the current needs are important compared to PIs from all other disciplines. Significantly more PIs from larger research groups (> five people) said seven of the current needs are important compared to those from smaller groups. Most of the differences between bioinformaticians and larger research groups persisted in their predictions of future needs (Table 1).

**Figure 2.**
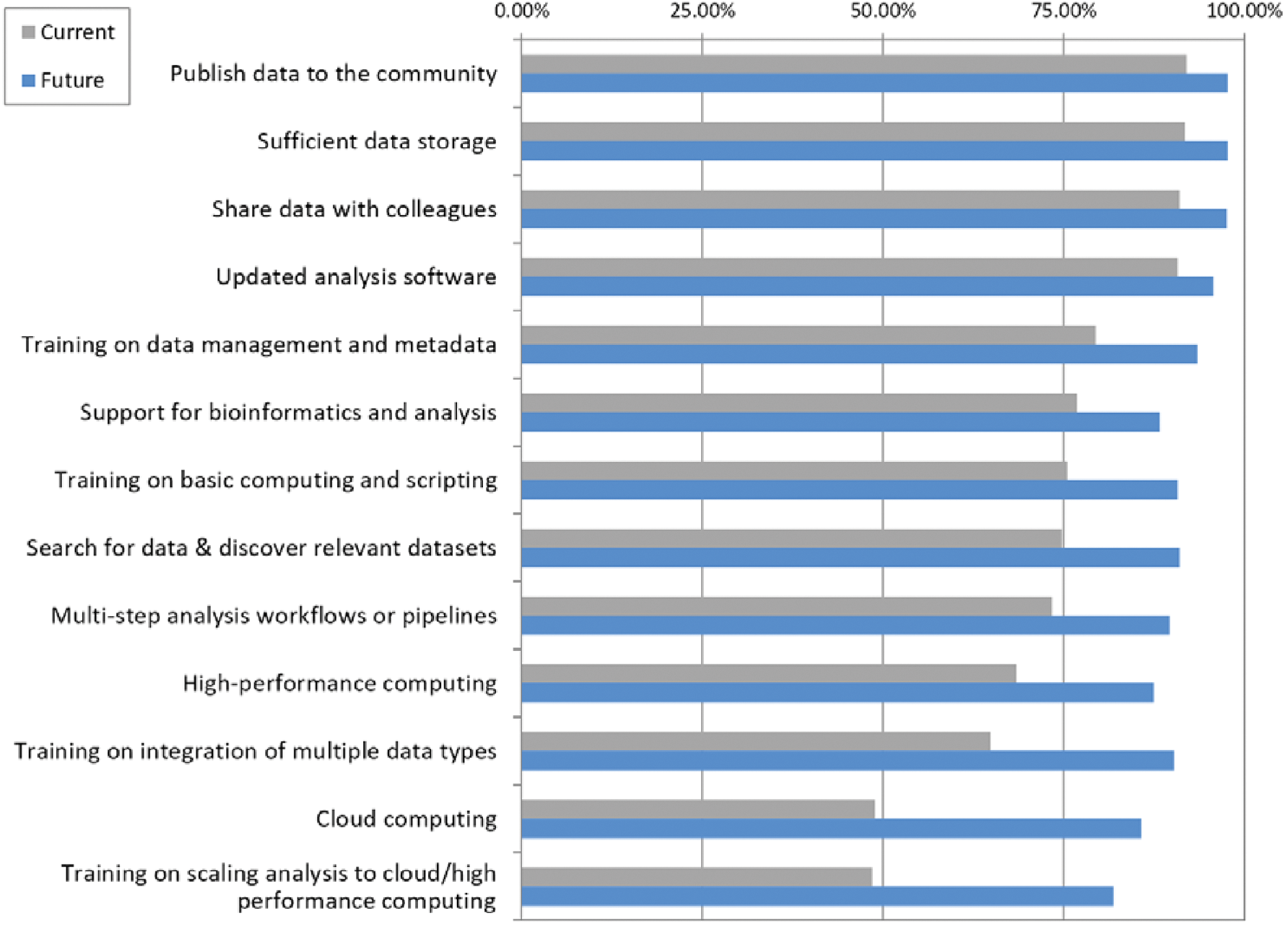
Percent of BIO PIs responding affirmatively that items are current (grey) or future (blue) computational needs. 387 ≤ n ≤ 551.

**Table 1.**
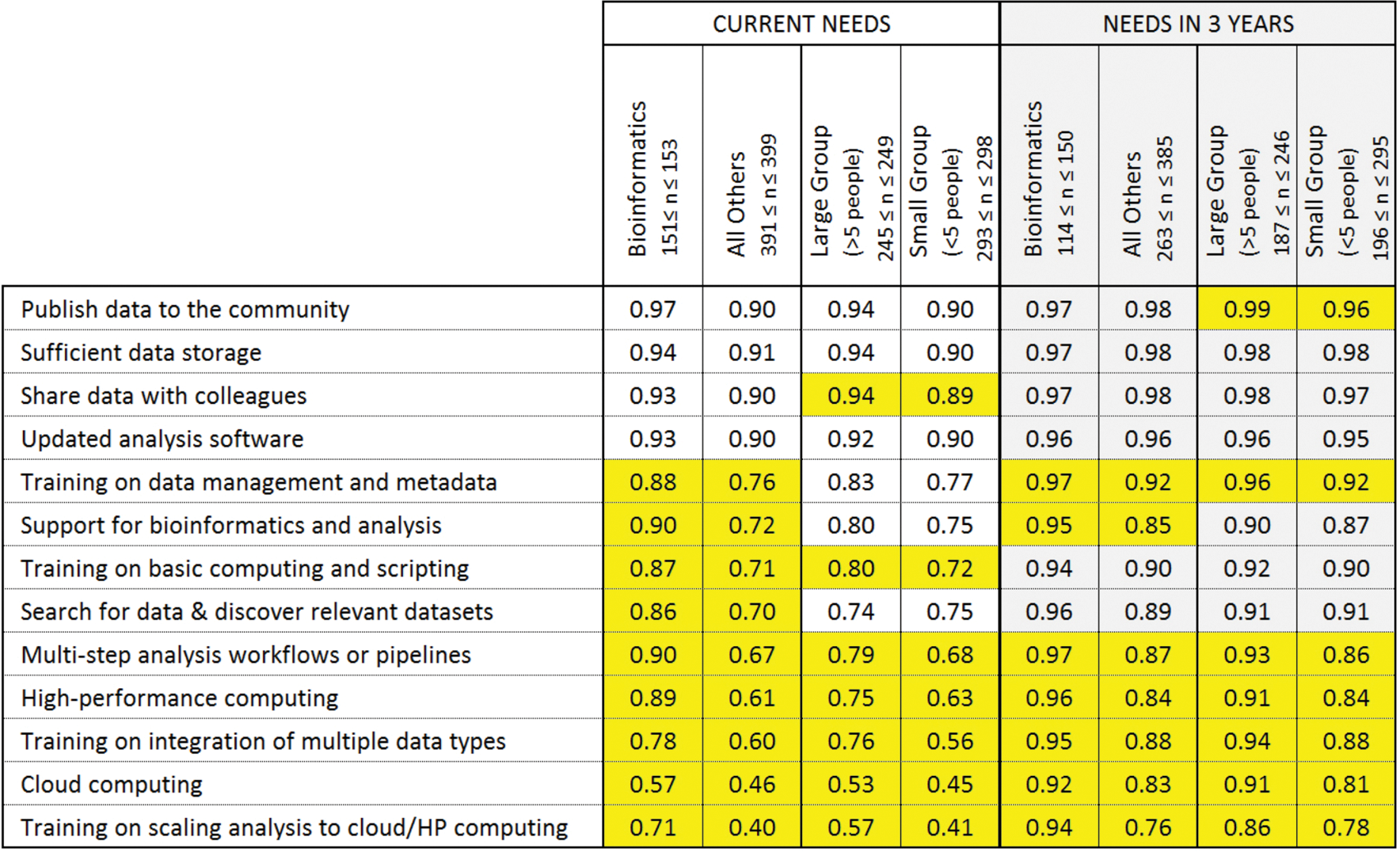
Percentage of affirmative responses, current and future data needs of BIO PIs. Yellow highlighting indicates a statistically significant chi square result between groups (bioinformaticians *versus* others; large research groups *versus* small).

Significantly more PIs funded by DEB said five of the current needs are important compared to PIs funded through the other three NSF research divisions. However, differences between the four NSF divisions disappeared for predictions of future need—suggesting that computational needs will converge across all fields of biology in the future (Table 2).

**Table 2.** Percentage of affirmative responses, current and future data needs of PIs, broken down by NSF BIO division. Yellow highlighting indicates a statistically significant chi square result between groups.

|  | CURRENT NEEDS |  |  |  | NEEDS IN 3 YEARS |  |  |  |
| --- | --- | --- | --- | --- | --- | --- | --- | --- |
|  | Environmental Biology<br>163 ≤ n ≤ 168 | Molecular and<br>Cellular Biosciences<br>85 ≤ n ≤ 90 | Biological Infrastructure<br>116 ≤ n ≤ 118 | Integrative Organismal<br>Systems<br>159 ≤ n ≤ 161 | Environmental Biology<br>124 ≤ n ≤ 162 | Molecular and<br>Cellular Biosciences<br>59 ≤ n ≤ 85 | Biological Infrastructure<br>85 ≤ n ≤ 117 | Integrative Organismal<br>Systems<br>108 ≤ n ≤ 157 |
| Publish data to the community | 0.95 | 0.92 | 0.93 | 0.87 | 0.99 | 0.98 | 0.96 | 0.98 |
| Sufficient data storage | 0.93 | 0.91 | 0.90 | 0.94 | 0.99 | 0.95 | 0.97 | 0.98 |
| Share data with colleagues | 0.95 | 0.90 | 0.91 | 0.88 | 0.98 | 0.99 | 0.96 | 0.97 |
| Updated analysis software | 0.92 | 0.88 | 0.91 | 0.91 | 0.95 | 0.93 | 0.95 | 0.99 |
| Training on data management and metadata | 0.87 | 0.71 | 0.81 | 0.74 | 0.94 | 0.91 | 0.95 | 0.92 |
| Support for bioinformatics and analysis | 0.80 | 0.83 | 0.72 | 0.76 | 0.89 | 0.90 | 0.88 | 0.87 |
| Training on basic computing and scripting | 0.83 | 0.77 | 0.65 | 0.72 | 0.94 | 0.89 | 0.85 | 0.92 |
| Search for data & discover relevant datasets | 0.75 | 0.77 | 0.77 | 0.71 | 0.93 | 0.93 | 0.91 | 0.88 |
| Multi-step analysis workflows or pipelines | 0.81 | 0.74 | 0.68 | 0.69 | 0.93 | 0.88 | 0.92 | 0.86 |
| High-performance computing | 0.77 | 0.68 | 0.64 | 0.63 | 0.91 | 0.90 | 0.85 | 0.83 |
| Training on integration of multiple data types | 0.69 | 0.57 | 0.62 | 0.65 | 0.91 | 0.93 | 0.89 | 0.91 |
| Cloud computing | 0.56 | 0.41 | 0.50 | 0.46 | 0.87 | 0.85 | 0.84 | 0.87 |
| Training on scaling analysis to cloud/HP computing | 0.55 | 0.46 | 0.50 | 0.41 | 0.86 | 0.78 | 0.79 | 0.80 |

A majority of PIs—across bioinformatics/other disciplines, larger/smaller groups, and the four NSF programs—said their institutions are not meeting nine of 13 needs (Figure 3). Training on integration of multiple data types (89%), on data management and metadata (78%), and on scaling analysis to cloud/HP computing (71%) were the three greatest unmet needs. High performance computing was an unmet need for only 27% of PIs—with similar percentages across disciplines, different sized groups, and NSF programs.

**Figure 3.**
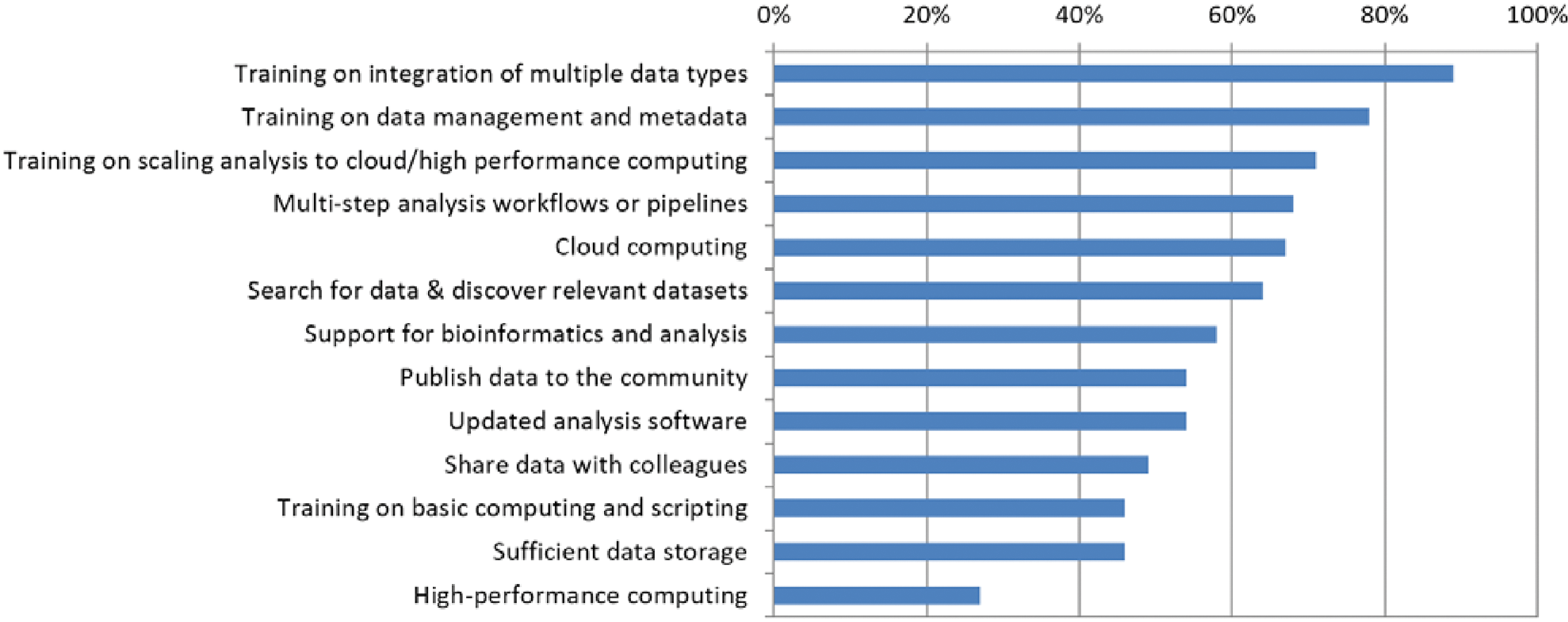
Unmet data needs of BIO PIs. Percent responding negatively (318 ≤ n ≤ 510).

## Discussion

This study fills a gap in the published literature on the computational needs of biological science researchers. Respondents had all been awarded at least one peer-reviewed grant from the BIO Directorate of NSF and thus represent competitive researchers across a range of biological disciplines. Even so, a majority of this diverse group of successful biologists did not feel that their institutions are meeting their needs for tackling large data sets.

This study stands in stark contrast to previous work, including one with a very different audience, which identified infrastructure and data storage as the most pressing computational needs. BIO PIs ranked availability of data storage and HPC lowest on their list of unmet needs. This provides strong evidence that the NSF and individual universities have succeeded in developing a broadly available infrastructure to support data-driven biology. Hardware is not the issue. The problem is the growing gap between the accumulation of many kinds of data—and researchers’ knowledge about how to use it effectively. The biologists in this study see training as the most important factor limiting their ability to best use the big data generated by their research.

Closing this growing data knowledge gap in biology demands a concerted effort by individual biologists, by institutions, and by funding agencies. We need to be creative in scaling up computational training to reach large numbers of biologists and in measuring the impact of our educational investments. Metrics for a super computer are readily described in terms of petaflops and CPUs, and we can facilely measure training attendance and “satisfaction.” However, answering unmet training needs will require a better understanding of how faculty are attempting to meet these needs—and how we can best assess their outcomes (*11*). For example, datasets available at the SRA provide almost unlimited entry points for course-based undergraduate research experiences (CUREs), which scale up discovery research in the context of for-credit courses. Participation in CUREs significantly improves student graduation rates and retention in science–effects that persist across racial and socioeconomic status (*12, 13*). However, many biologists acquire skills for big data analysis on their own. Software Carpentry and Data Carpentry (*14*) are volunteer-driven organizations that provide a cost-effective, disseminated model for reaching biologists outside of an academic classroom.

Reflected in the top two unmet needs of BIO PIs is the looming problem of integrating data from different kinds of experiments and computational platforms. This will be required for a deeper understanding of “the rules of life” (*15, 16*)—notably the genotype-environment-phenotype interactions that are essential to predicting how agricultural plants and animals can adapt to changing climates. Such integration demands new standards of data management and attention to metadata about how these data are collected. The BIO PIs in this study are anticipating a new world of pervasive data in which they will need to become data scientists.

## Acknowledgements

This study is an Education, Outreach and Training (EOT) activity of CyVerse, an NSF-funded project to develop a “cyber universe” to support life sciences research (DBI-0735191 and DBI-1265383). The authors also wish to thank Bob Freeman and Christina Koch of the ACI-REF project for helpful discussions and references during the development of the survey.

## Data availability

Data are available for download at: https://doi.org/10.6084/m9.figshare.4643641

